# A Systematic Evaluation of Protein Phase Separation Predictors Across Diverse Protein Landscapes

**DOI:** 10.64898/2026.01.20.700394

**Authors:** Kate E Gilroy, John N Barr

## Abstract

**Background:** Liquid–liquid phase separation (LLPS) plays a central role in cellular regulation, with its dysregulation linked to numerous disease outcomes. LLPS has also being increasingly implicated as critical in various biological contexts, with a key example being virus replication. These findings have driven the development of numerous computational predictors to screen and identify phase-separating proteins from sequence and/or structural models. Despite the growing need for these tools, their comparative performance across diverse biological contexts remains incompletely understood, complicating tool selection and result interpretation.

**Results:** A systematic comparative analysis of nine LLPS prediction algorithms was conducted using multiple curated datasets comprising both LLPS-positive (LLPS+) and -negative (LLPS-) proteins. The datasets span a range of biologically relevant scenarios, including intrinsically disordered proteins, folded proteins, proteins with LLPS-abolishing variants, benchmark datasets, and viral proteins. Substantial variability in predictive performance was observed across algorithms when assessing proteins of different structural classes. Numerous showed reduced accuracy in distinguishing LLPS+ and LLPS- folded proteins. Sensitivity to the impact of small LLPS-abolishing mutations also varied markedly between predictors, with structure-informed algorithms generally outperforming most sequence-based predictors. Similar decreases in predictive capability were also observed across several algorithms when analysing viral proteins, which are often-under-represented in predictor training datasets.

**Conclusions:** These results demonstrate that LLPS predictor performance is strongly context dependent, leading to different predictors being optimal for different biological questions. For overall protein assessment, DeePhase and Molphase provided the most consistently accurate predictions, being the least impacted by structural bias. For assessing the impact of small mutations on LLPS propensity, PSPHunter, a built-for-purpose algorithm, reliably predicts mutation impacts with structure-informed algorithms PSPire and PICNIC also providing strong insight. Across all evaluated datasets, the findings highlight the need for well-benchmarked training and testing data that encompasses a broad and diverse range of protein classes.

## Background

Biomolecular condensates (BMCs) are cellular organelles or structures that lack a delimiting membrane but still function in concentrating and segregating cellular components (1–4). In recent years the regulation of such structures has been identified as often occurring via the process of liquid-liquid phase separation (LLPS), causing the demixing of macromolecular components in a solution (5). This finding has led to greater understanding of numerous essential cellular structures and processes, including P-bodies and stress granules (2,6), and its dysregulation has been linked to numerous disease outcomes (7–11). While there have been great strides made in the ability to experimentally identify phase-separating proteins (PSPs), it remains a laborious task often requiring extensive *in vitro* and *in cellulo* investigations (12). Due to the increased recognition of the importance of LLPS in the regulation and dysregulation of biological systems, more high-throughput screening methods are essential to identify potential PSPs.

Some of the earliest computational predictors used for LLPS date back to 2014 (13) but were not initially designed for specific prediction purposes. PLAAC predicts prion-like domains (PLDs) (13), R+Y assesses the arginine and tyrosine content of a protein (14), and PScore calculates planar pi-interaction frequency (15). After their production, however, researchers recognised their utility as LLPS predictors, since many of the characteristics these algorithms assess are identifiers of PSPs. Despite their importance early on, these methods have now become out-dated by a second-generation of designed-for-purpose LLPS prediction algorithms, many of which incorporate machine learning models, learning patterns from a wide array of features including amino acid composition and intrinsic disorder, rather than the single-feature approach. These algorithms also have access to a larger range of datasets for testing and training, increasing their abilities.

The majority of predictors look at “sequence-based” features, focussing on biochemical properties of each amino acid within the primary protein sequence, as it has been proposed this is where the LLPS-capacity of a protein is defined (16). However, it has been argued that sequence features alone are insufficient for accurate prediction, and structural aspects also need to also be considered (17). Adding structural features, however, had been limited due to the lack of high-quality protein structures. However, since the wide availability of Alphafold (18,19) to the public, this has changed, with near-experimental quality structures being produced and deposited in the Alphafold Protein Structure Database (20,21). This has led to the recent development of a potential third-generation of LLPS predictors that assess not only sequence but also consider structure-based characteristics such as structured superficial regions (SSUPs) and secondary arrangements. Predictors employing this third-generation approach include PICNIC (22) and PSPire (17), claiming to identify PSPs with higher accuracy than both first- and second-generation algorithms. Interestingly, both also assert the ability to identify PSPs regardless of intrinsically disordered regions (IDRs), one of the most well-established features driving LLPS (23). Due to the large contribution of IDRs to LLPS, most predictors consider it as a main indicator of phase separation potential, however, a growing number of proteins with no, or limited, IDRs have been found to undergo phase separation (17,24). Additionally, the mere presence of IDRs is not enough for many proteins to phase separate under physiological conditions (25) and thus this bias in predictors can be leading to both false positive and false negative results. This poses the question of whether third generation, structure-informed predictors that look beyond IDRs will provide the most accurate results or if structure is not required for accurate LLPS predictions.

Despite these recent advances and new avenues in LLPS prediction algorithms, there have been few unbiased comparative assessments of these different predictive approaches. Liao et al. (26) preformed a comparative analysis of numerous first- and second-generation algorithms, confirming the better predictive performance of machine learning-based approaches. However, in the few years since their study, the third-generation of structure-based, IDR-unbiased approaches have been released, requiring assessment and comparative analysis of predictor performance using structural models. There are also additional unaddressed concerns regarding the usefulness of these algorithms beyond some limited scenarios. These include biases in the training and testing datasets used to develop these predictors, the ability to detect changes in LLPS propensity caused by small mutations, the ability to identify regions responsible for driving phase separation, and the ability to accurately predict phase separation for underrepresented organisms with limited homologues, such as viruses (27–29).

To address these issues, here, we performed a comprehensive comparative analysis of nine second- and third-generation LLPS predictors: Molphase (30), Deephase (31), Fuzdrop (32–34), PSPHunter (35), LLPhyscore (16), PSAP (36), PSPire (17), Phaseek (37), and PICNIC (22). Each predictor was compared across a variety of datasets including a ‘standard’ testing dataset, a selection of LLPS-abolishing mutants, and a set of LLPS+ and LLPS- viral protein sequences. Additionally, the effectiveness of third-generation predictors utilising structural protein models was compared to that of the sequence-based second-generation predictors. The performance of all algorithms was then analysed for overall prediction power in addition to accuracy in specialist scenarios to aid researchers in identifying the best performing algorithm for their applications.

## Methods

### Selecting Phase Separation Predictors for Evaluation

A targeted literature search was undertaken to identify second-generation phase separation predictors. This included those employing machine or deep learning algorithms, and/or those utilising multiple engineered features and large datasets specifically designed for phase separation prediction. The search was then extended to include predictors that assess structural protein models in addition to sequence features – the potential third-generation algorithms. This returned 11 predictors, eight second generation – Molphase (30), DeePhase (31), Fuzdrop (32–34), PSPHunter (35), Seq2Phase (38), PSPredictor (39), LLPhyScore (16), PSAP (36), and Phaseek (37) – and two third generation - PSPire (17) and PICNIC (22). Of these predictors, two were eliminated: PSPredictor and Seq2Phase. PSPredictor was excluded as, at the time of this analysis, the web-based interface was unavailable, and no code was made publicly available to run locally, making it inaccessible. The other excluded predictor, Seq2Phase, provides the code required, however it involves the user training the Seq2Phase model locally. This requires a large amount of computing power to be completed quickly and successfully, making it inaccessible to many potential users. Thus, the focus of this study was placed on algorithms that are most accessible for researchers of all backgrounds. Key details of the nine selected algorithms can be seen in Table 1.

**Table 1.**
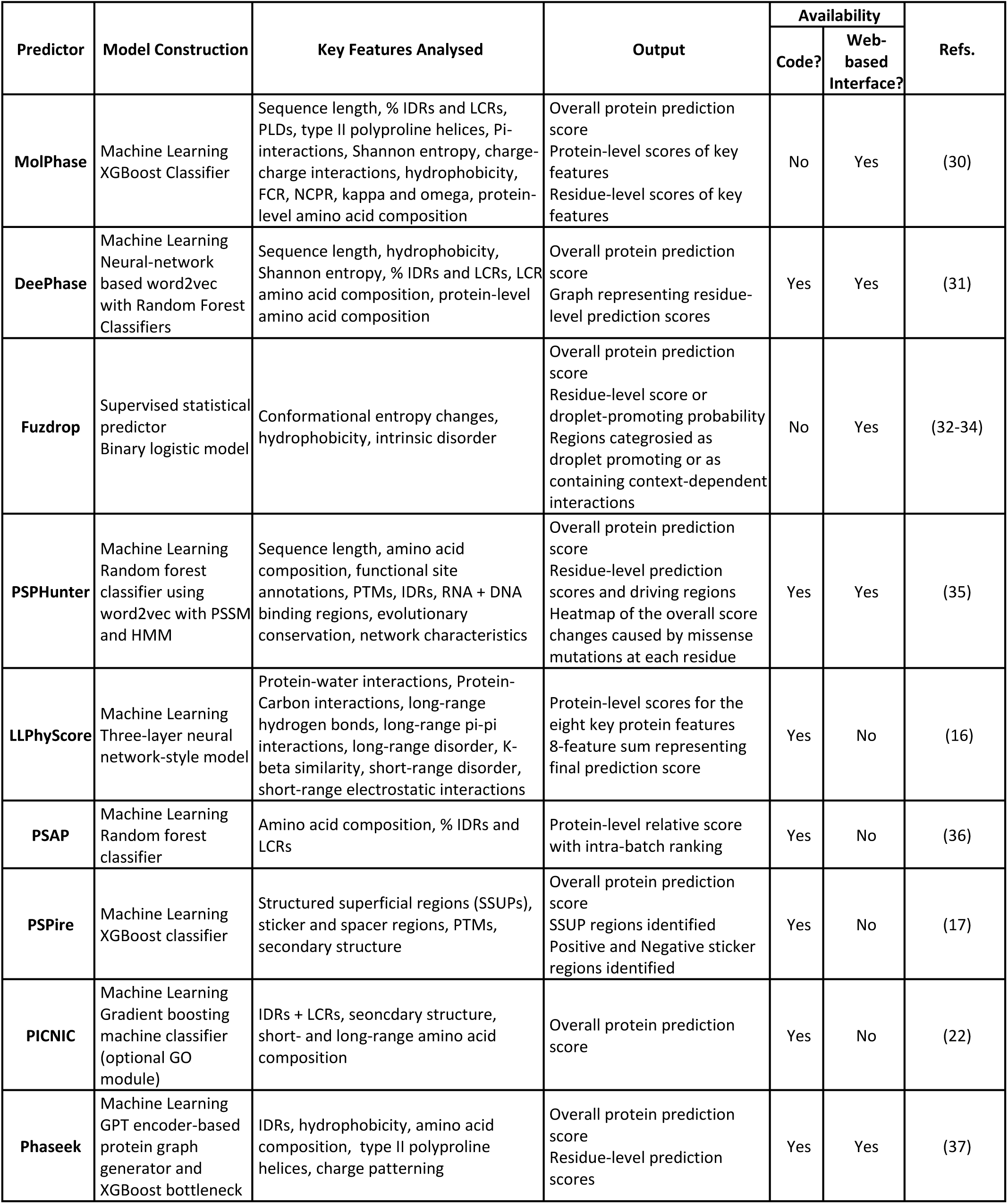
Key Details of Chosen LLPS Predictors.

| Predictor | Model Construction | Key Features Analysed | Output | Availability |  | Refs. |
| --- | --- | --- | --- | --- | --- | --- |
|  |  |  |  | Code? | Web-based Interface? |  |
| **MolPhase** | Machine Learning XGBoost Classifier | Sequence length, % IDRs and LCRs, PLDs, type II polyproline helices, Pi-interactions, Shannon entropy, charge-charge interactions, hydrophobicity, FCR, NCPR, kappa and omega, protein-level amino acid composition | Overall protein prediction score<br>Protein-level scores of key features<br>Residue-level scores of key features | No | Yes | (30) |
| **DeePhase** | Machine Learning Neural-network based word2vec with Random Forest Classifiers | Sequence length, hydrophobicity, Shannon entropy, % IDRs and LCRs, LCR amino acid composition, protein-level amino acid composition | Overall protein prediction score<br>Graph representing residue-level prediction scores | Yes | Yes | (31) |
| **Fuzdrop** | Supervised statistical predictor<br>Binary logistic model | Conformational entropy changes, hydrophobicity, intrinsic disorder | Overall protein prediction score<br>Residue-level score or droplet-promoting probability<br>Regions categorised as droplet promoting or as containing context-dependent interactions | No | Yes | (32-34) |
| **PSPHunter** | Machine Learning Random forest classifier using word2vec with PSSM and HMM | Sequence length, amino acid composition, functional site annotations, PTMs, IDRs, RNA + DNA binding regions, evolutionary conservation, network characteristics | Overall protein prediction score<br>Residue-level prediction scores and driving regions<br>Heatmap of the overall score changes caused by missense mutations at each residue | Yes | Yes | (35) |
| **LLPhyScore** | Machine Learning Three-layer neural network-style model | Protein-water interactions, Protein-Carbon interactions, long-range hydrogen bonds, long-range pi-pi interactions, long-range disorder, K-beta similarity, short-range disorder, short-range electrostatic interactions | Protein-level scores for the eight key protein features<br>8-feature sum representing final prediction score | Yes | No | (16) |
| **PSAP** | Machine Learning Random forest classifier | Amino acid composition, % IDRs and LCRs | Protein-level relative score with intra-batch ranking | Yes | No | (36) |
| **PSPire** | Machine Learning XGBoost classifier | Structured superficial regions (SSUPs), sticker and spacer regions, PTMs, secondary structure | Overall protein prediction score<br>SSUP regions identified<br>Positive and Negative sticker regions identified | Yes | No | (17) |
| **PICNIC** | Machine Learning Gradient boosting machine classifier (optional GO module) | IDRs + LCRs, seoncdary structure, short- and long-range amino acid composition | Overall protein prediction score | Yes | No | (22) |
| **Phaseek** | Machine Learning GPT encoder-based protein graph generator and XGBoost bottleneck | IDRs, hydrophobicity, amino acid composition, type II polyproline helices, charge patterning | Overall protein prediction score<br>Residue-level prediction scores | Yes | Yes | (37) |

### Dataset Construction

To avoid any overlap of sequences used in this analysis to those used to train the predictors, all training sequences from each predictor were collected from the Methods or Supplementary Materials section of each manuscript. For each dataset described below, proteins that had more than 40% sequence identity with any training sequence were identified and removed via CD-HIT (40).

#### Standard Dataset

The standard dataset was created to represent the commonly used process of generating positive and negative datasets for both training and testing most prediction algorithms. For the LLPS-positive (LLPS+) dataset, driver-exclusive, experimentally validated PSP sequences were collected from the database MLOsMetaDB (41). For the LLPS-negative (LLPS-) dataset, the Protein Sequence Culling Server (PISCES) (42) was used to collect protein sequences with high-resolution X-ray structures containing no chain breaks or breaks due to disorder. These proteins are commonly used as negative sequences as those with low-to-no disorder and well-established structures are unlikely to undergo phase separation. Any proteins containing non-canonical amino acids, shorter than 60 amino acids, or with over 40% sequence identity between positive and negative classes were removed. Biopython SeqIO (43) was then utilised to randomly select 100 positive and 100 negative sequences, forming the *Standard* dataset.

#### Disordered vs. Folded Dataset

Despite the evidence that proteins with ordered structure are less likely to phase separate, and those with disorder are more likely, this is not always true (17,24). To assess if the predictors are affected by this bias, LLPS+ and LLPS- classes were further categorised into ‘disordered’ or ‘folded’. The LLPS+ sequences were generated as per the *Standard* dataset, however prior to condensing to 100 sequences, disorder was assessed via MetaPredict (44). Sequences with MetaPredict scores <0.5 were considered ‘folded’ and those with scores ≥0.5 were considered ‘disordered’. SeqIO was then utilised to collect 100 randomised sequences of each.

The LLPS- dataset outlined in the *Standard* dataset was utilised as the ‘folded’ category with another 100 sequences being randomly selected by SeqIO. For the LLPS- ‘disordered’ proteins, the database DisProt (45) was used. All non-obsolete sequences were collected and those marked as ‘condensates-related proteins’ by DisProt were removed. Additionally, any sequences with >40% sequence similarity to any folded or disordered proteins in this dataset were removed by CD-HIT, as well as those with non-canonical amino acids or sequences shorter than 60 residues. 100 sequences were chosen at random by SeqIO.

#### Benchmark Dataset

It is currently hypothesised that almost any protein, regardless of structural status, has the potential to undergo phase separation under specific, though often non-physiological, conditions (46,47). This makes it difficult to produce a ‘true negative’ dataset for predictor training and testing as there are almost certainly false negatives included. To challenge this, Pintado-Grima et al. (2025) have attempted to produce high-confidence ‘benchmark’ LLPS+ and LLPS- datasets utilising strict filters for curation. For the LLPS+ dataset, proteins identified as ‘driver-exclusive’ were selected, and the full LLPS- dataset was used. Any proteins containing non-canonical amino acids, shorter than 60 amino acids, or with over 40% sequence identity between positive and negative classes were removed. SeqIO was then utilised to randomly select 100 positive and 100 negative sequences

#### Mutations Dataset

It has been well documented that in many proteins, small and/or single amino acid mutations can significantly alter or abolish the ability of a protein to undergo phase separation (48–50). It is therefore important to assess the capability for predictors to detect these small, but significant changes and reflect this in their scores. LLPSDB v2.0 (51) allows filtration of PSPs to identify those in which the wildtype (WT) sequence undergoes LLPS, but specific mutations have been experimentally validated to disrupt this ability. Following culling of training proteins, 12 WT sequences remained. Unique mutation sequences from LLPSDB v2.0 for the corresponding WT sequences were then collated. In this dataset the LLPS+ proteins are the WT sequences and the LLPS- are the mutant sequences.

#### Virus Dataset

To assess if the predictors can accurately assess the LLPS ability of proteins from an under-represented class, viral proteins were collected. LLPS+ viral sequences were collected from BAV-LLPS-DB (52) and unsuitable proteins filtered out as with previous datasets, resulting in 30 LLPS+ viral sequences.

For LLPS-, viral sequences with high-resolution X-ray structures were collected from PDB and filtered as with previous datasets. 30 random sequences were then selected by SeqIO.

As PSPire and PICNIC require structural files for their predictions, with a preference for AlphaFold structures, the corresponding structures were collected from the AlphaFold database (20,21). However, where there was no structure available, AlphaFold2 (18,19) was used to produce batch predictions. An overview of the dataset construction workflow can be seen in Figure 1 and the sequences used in each dataset are outlined in Additional File 1.

**Figure 1.**
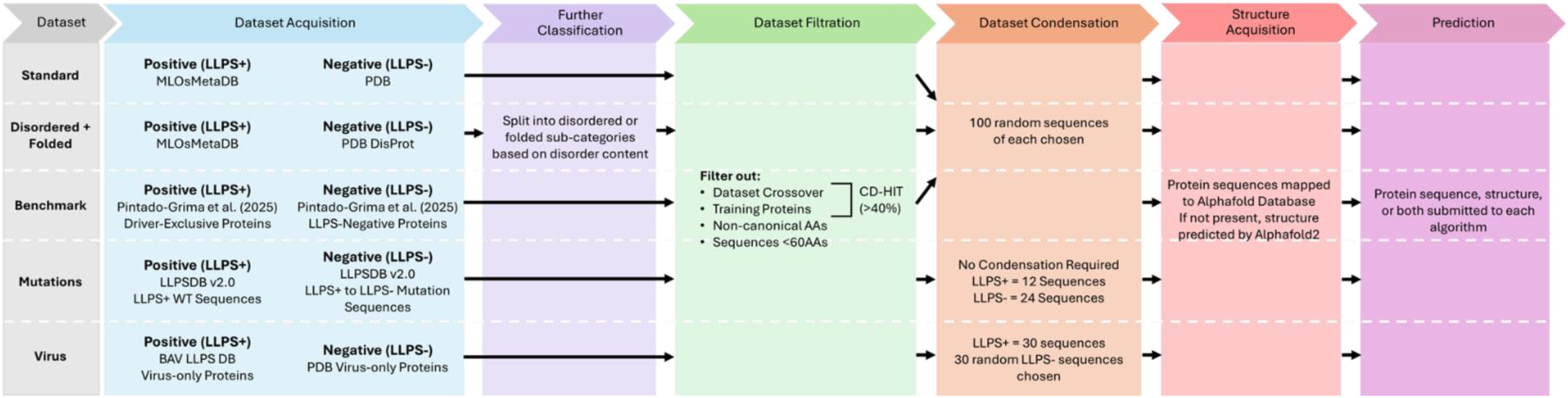
Workflow of Dataset Construction. Overview of the acquisition, filtration, and analysis steps taken for each dataset utilised in this study.

### Data Analysis and Evaluation Parameters

To assess the overall performance and accuracy of the predictors, receiver operating characteristic (ROC) curves and precision-recall curves (PRC) were constructed. Area under the curve for each was calculated (AUROC and AUPRC, respectively) utilising python package sklearn (53).

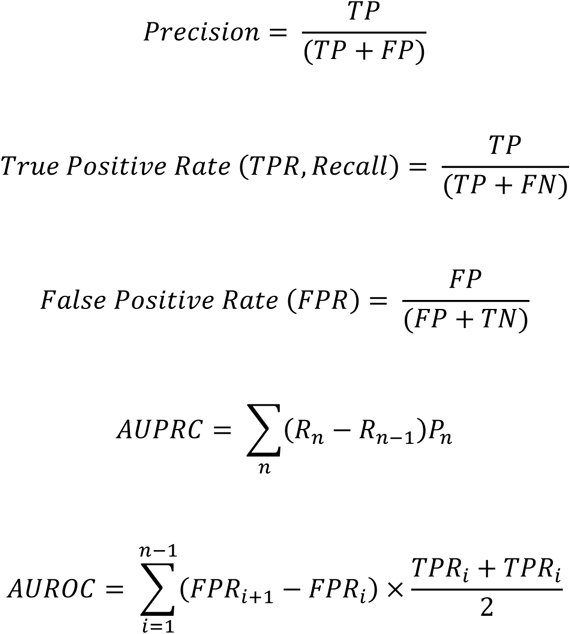

In the above equations: TP is true positive, FP is false positive, FN is false negative, TN is true negative, R is recall, P is precision, n is threshold, and i is index.

To calculate fold change in prediction score with the *Mutations* dataset the following equation was used:

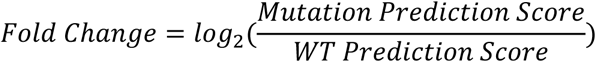

For generating a final ‘rank’, each algorithm was scored from 1 to 9 on performance in each individual dataset with the highest AUROC score being rank 1 and the lowest AUROC score being rank 9. The modal rank for each algorithm across all datasets was then calculated and returned as the overall rank.

## Results and Discussion

### Performance of Prediction Algorithms on a Standard LLPS Testing Dataset

In the production, training, and testing of LLPS algorithms, there are common methods in which datasets are curated. For LLPS+ proteins, curation has been made simpler via the recent advances in identifying these proteins experimentally and the subsequent development of databases to collate and categorise them (41,54–57). For LLPS- datasets it is significantly more complex. Due to the nature of LLPS, it is near impossible to identify a protein as a ‘true negative’ as many proteins only undergo LLPS under specific conditions (12,58), of which it would not be feasible to test every protein against. Instead, negative datasets are often built based on the idea that proteins with well-documented non-flexible structures are much less likely to undergo phase separation than those with a flexible, disordered nature. Therefore, taking a collection of proteins with high-resolution X-ray structures from databases like PDB (59) is the most common method of building a LLPS-dataset.To test these algorithms with a ‘native’ dataset, i.e. one similar to what they were trained and tested with initially, the *Standard* dataset was produced. Experimentally validated LLPS+ proteins were collected from an array of different LLPS+ databases via the meta-database MLOsMetaDB (41) and LLPS- proteins were identified as those with high-resolution X-ray structures from PDB.

To assess the performance of each predictor, receiver operator characteristic (ROC) curves and precision recall curves (PRC) were constructed and the area under value was then calculated: AUROC and AUPRC respectively. Despite all predictors being trained and tested on similarly curated datasets, only five of the nine algorithms possess AUROC scores with very high accuracy (>0.9) (Fig. 2a). Interestingly, the two structure-informed predictors included in this analysis had a good level of accuracy (PSPire: 0.741, PICNIC: 0.682) but were middling in comparison to the others assessed (Fig. 2c). This suggests that while structural features are key for the process of LLPS, sequence-derived features alone are sufficient to provide highly reliable LLPS propensity prediction.

**Figure 2.**
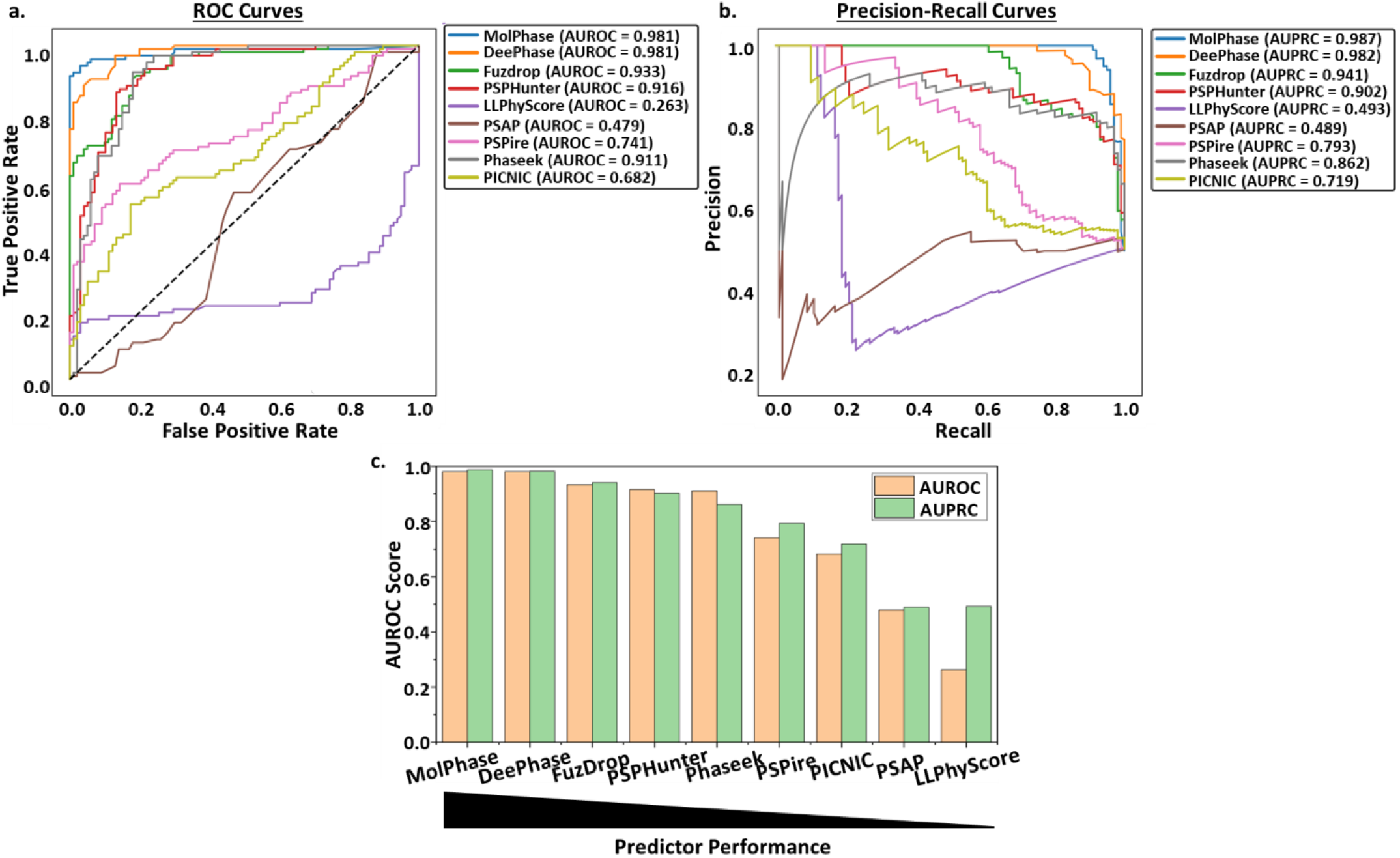
Analysis of LLPS predictors using a *Standard* dataset. **a.** ROC curves and **b.** PRCs of all nine algorithms following prediction of the *Standard* protein dataset. AUROC and AUPRC calculated for each. Black dashed line represents an AUROC of 0.5. **c.** Predictors ranked via AUROC performance from highest (left) to lowest (right).

Surprisingly, two predictors performed below random chance (0.5), actively placing proteins into the opposing categories: PSAP (0.479) and LLPhyScore (0.263). LLPhyScore produces a linear weighted sum score of the top 8 contributing sequence factors as the final propensity score. The score considers whether these features are positively (e.g. disorder) or negatively (e.g. short-range electrostatics) correlated to a higher LLPS propensity. While this does allow for an easier interpretability of which features contribute to the overall score and to what extent, it also leads to vulnerability in the predictions. Proteins undergoing LLPS can have very diverse characteristics, contributing to the struggle to find the key features for identification and prediction. Therefore, a protein that has been experimentally shown to phase separate but has an ‘outlier’ characteristic (e.g. low disorder or high electrostatics) could be incorrectly predicted by LLPhyScore. This appears to be less of an issue with other predictors, such as MolPhase and DeePhase, which utilise larger numbers of sequence properties and non-linear machine/deep learning (e.g. random forest) to better understand the complex relationships between characteristics and handle ‘outlier’ features. This does sacrifice some level of interpretability of each feature’s contribution but provides more accurate overall predictions. This also aligns well with the AUPRC score being much higher than the AUROC for LLPhyScore (Fig 2b,c), suggesting the proteins it is scoring correctly are only those with a high LLPS propensity.

PSAP scored much closer to the ‘random chance’ boundary but was still not performing as well with this dataset. While it does utilise a large number of protein sequence features and a non-linear random forest approach, issues likely come from the training. For this predictor, it was trained on only 90 high-confidence, experimentally identified PSPs. This is a relatively low number of proteins to use for training purposes as it can reduce the generalisability of the predictions, meaning proteins that do phase separate but have less ‘characteristic’ features (e.g. lower disorder, no low complexity regions, etc.) are more likely to be incorrectly assessed.

### Disordered and Folded Proteins Reveal Structural Bias in Phase Separation Prediction

One of the issues with using a dataset like the *Standard* set for training and testing predictors is that it limits the diversity of the proteins included. Many proteins that are disordered have been shown to undergo phase separation (23), however, not every disordered protein shows a ready propensity for this (23,60,61). By using highly ordered protein sequences from the PDB as the main source of LLPS- proteins, the predictions made will therefore be biased towards ordered proteins being negative and disordered proteins being positive, which is not always correct. To assess if the predictors are impacted by this bias, the *Disordered vs. Folded* dataset was produced. As shown in Fig. 1, LLPS+ proteins were collated from MLOsMetaDB and categorised into disordered or folded utilising MetaPredict (44) (threshold of 0.5). LLPS- folded proteins were collected as usual from the PDB but the disordered LLPS- proteins were taken from the DisProt (45) disordered protein database.

The mixed dataset (Fig. 3a) is similar in composition to the *Standard* dataset (Fig. 2) but is equally balanced with folded and disordered proteins of both LLPS+ and LLPS- classification. While the results initially seem similar to those of the *Standard* dataset, six of the nine predictors showed a decrease in overall AUROC and AUPRC scores (Fig. 3ai,ii). This suggests a potential over-fitting of the predictors to the idea that disordered means LLPS+ and folded means LLPS- as the more diverse the test set, the weaker the scores. Those that showed slight increases in AUROC and AUPRC (PSPire, PSAP, and LLPhyScore) still, however, remain middling-to-low in overall comparative performance (Fig. 3aiii).

**Figure 3.**
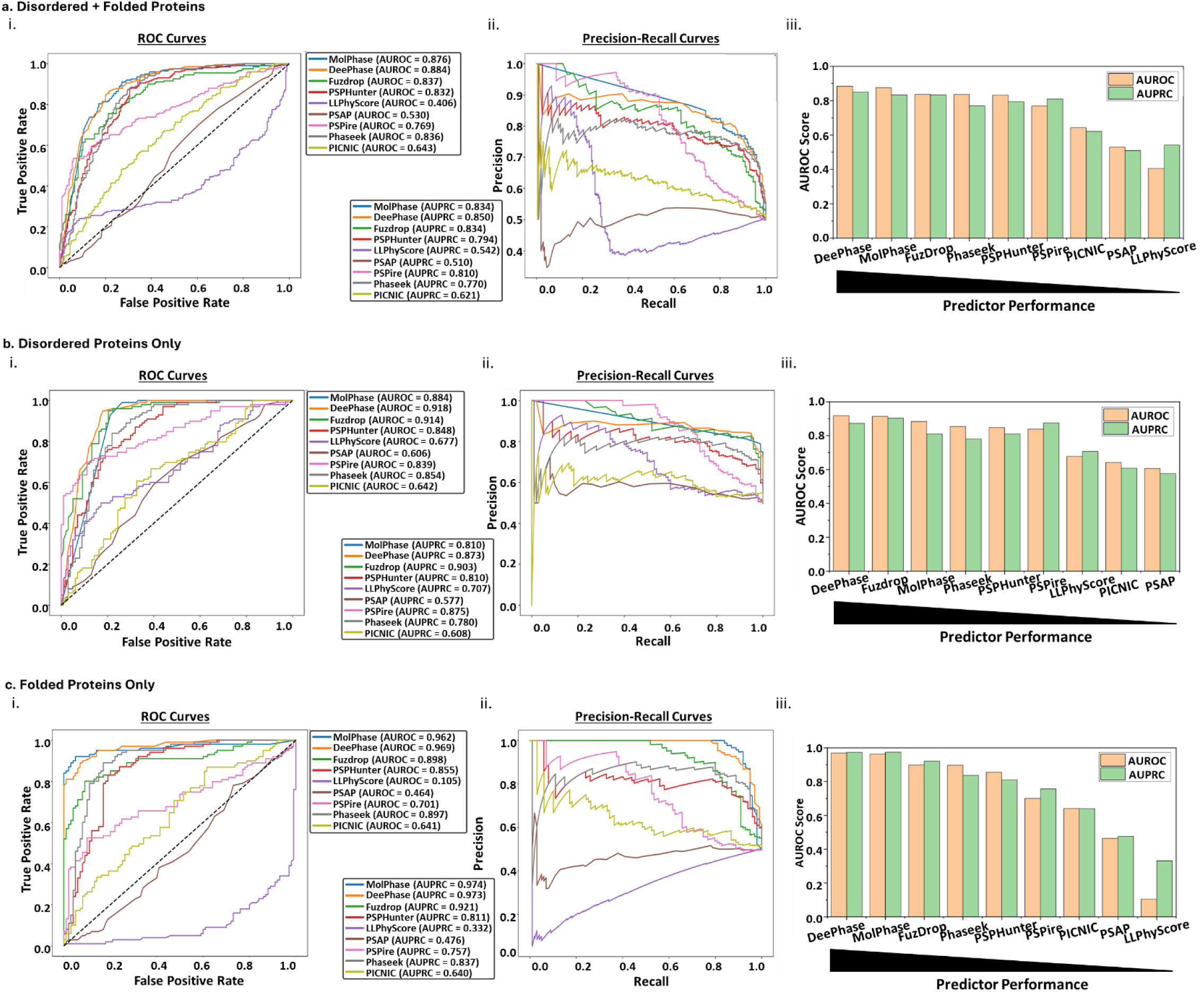
LLPS predictors vary in the ability to discriminate LLPS+ and LLPS- proteins of different structural order. The performance of each LLPS predictor was assessed on datasets consisting of equally mixed LLPS+ and LLPS- proteins. The datasets contained either **a.** an equal mixture of disordered and folded proteins, **b.** only disordered proteins, or **c.** only folded/structured proteins. For each dataset the **i.** AUROC and **ii.** AUPRC were calculated for each predictor which were then **iii.** ranked based on AUROC score.

To further understand how these biases could be affecting the predictors, the different structure types were run individually through each algorithm. As expected, when assessing only disordered LLPS+ and LLPS-proteins, the same trend of decreased AUROC and AUPRC as with the mixed set is evident in the same six predictors (Fig 3bi,ii). This solidifies the idea that many of these algorithms are over-fitted, likely due to the over-reliance on structured proteins for a negative dataset. Interestingly, one of the only algorithms showing an increase in AUROC and AUPRC in this dataset is PSPire, a structure-informed algorithm. PSPire was specifically designed to make assessments based on protein structure files while also not relying heavily on disorder for its predictions (17), something reflected by this dataset. The other two predictors showing an increase were the two lowest performers with the S*tandard* dataset: LLPhyScore and PSAP. Both now score >0.6, well above the random chance line which indicates that, unlike the other algorithms, these are not overfitted to the same idea of disorder.

However, the issues in LLPhyScore’s model becomes evident when considering only folded LLPS+ and LLPS-proteins (Fig. 3c). The AUROC score from the disordered only dataset decreased by 0.572 when looking at folded proteins, suggesting that LLPhyScore is unable to discriminate the LLPS propensity of a well-structured protein, relying heavily on disorder attributes. In comparison to the standard dataset, all algorithms saw a decrease in AUROC and AUPRC scores, though most were only small changes.

Overall, dissection of predictor performance based on structural disorder of a protein has highlighted the common overfitting of predictors to the idea that the higher the level of disorder, the higher the propensity for LLPS (and vice versa) when this is becoming less of a scientifically accepted idea. Therefore, it is a necessity to develop more diverse training datasets to produce these predictors and aid in avoidance of this disorder bias. Promisingly, however, the same three algorithms have formed the top performers across the *Standard* (Fig. 2c) and *Disorder vs. Folded* (Fig. 3aiii, biii, ciii) datasets – DeePhase, MolPhase, and FuzDrop. While being somewhat affected by bias in training, each of these predictors showed high levels of accuracy (AUROC and AUPRC >0.8) no matter the dataset, suggesting these to be highly robust predictors for diverse proteins.

### Benchmarking Predictors on a Diverse and High-Confidence Dataset Uncovers Performance Disparities

As previously discussed, a large confounding factor in the prediction of LLPS proteins is the lack of standardised benchmark datasets, especially for LLPS- proteins. The issues with this have been exemplified here with poorer prediction scores being returned depending on the composition and diversity of the datasets to be predicted. To tackle this Pintado-Grima et al. (62) have developed more strictly filtered LLPS+ and LLPS- sets of proteins from a wide variety of LLPS databases and protein structure sites, including DisProt for disordered LLPS- representation. Part of this strict filtration includes the removal of any LLPS- proteins with documented interactions with any known PSPs or proteins related to LLPS. This should help reduce the inclusion of false negatives while retaining a higher level of structural diversity than standard datasets.

While the effects of using this *Benchmark* dataset for training a predictor will not be tested here, utilising it as a testing dataset can give some insight into suitability of the current training of these predictors. For example, when compared to the results from the *Standard* dataset, if the AUROC score increases, it suggests the algorithm performs well and was being ‘dragged down’ by incorrectly labelled proteins, most likely false negatives. In the opposite case with a decrease in AUROC, it suggests that the predictors may have learned incorrect correlations or artefacts from being trained on ‘noisy’ data.

The top four performers from the previous datasets (MolPhase, DeePhase, Fuzdrop, and Phaseek) all show a ∼0.1 decrease in AUROC compared to the *Standard* dataset (Fig. 4a), suggesting impacts of ‘noisy’ training data. MolPhase, DeePhase, and Phaseek all employ only PDB sequences as the negative dataset, likely accounting for these issues. Interestingly, with DeePhase there is a large difference between the AUROC and AUPRC (Fig. 4b). While the AUROC decreased, the AUPRC stayed near *Standard* dataset levels, suggesting that DeePhase still performs well with identifying true positives and thus the issues are likely with prediction of the negative class. Fuzdrop, on the other hand, employs the human proteome (minus known PSPs) as its negative dataset. While this does improve upon the PDB-based datasets in terms of structural and protein diversity, it also very likely contains many false negatives that are yet to be identified, contributing to noise.

**Figure 4.**
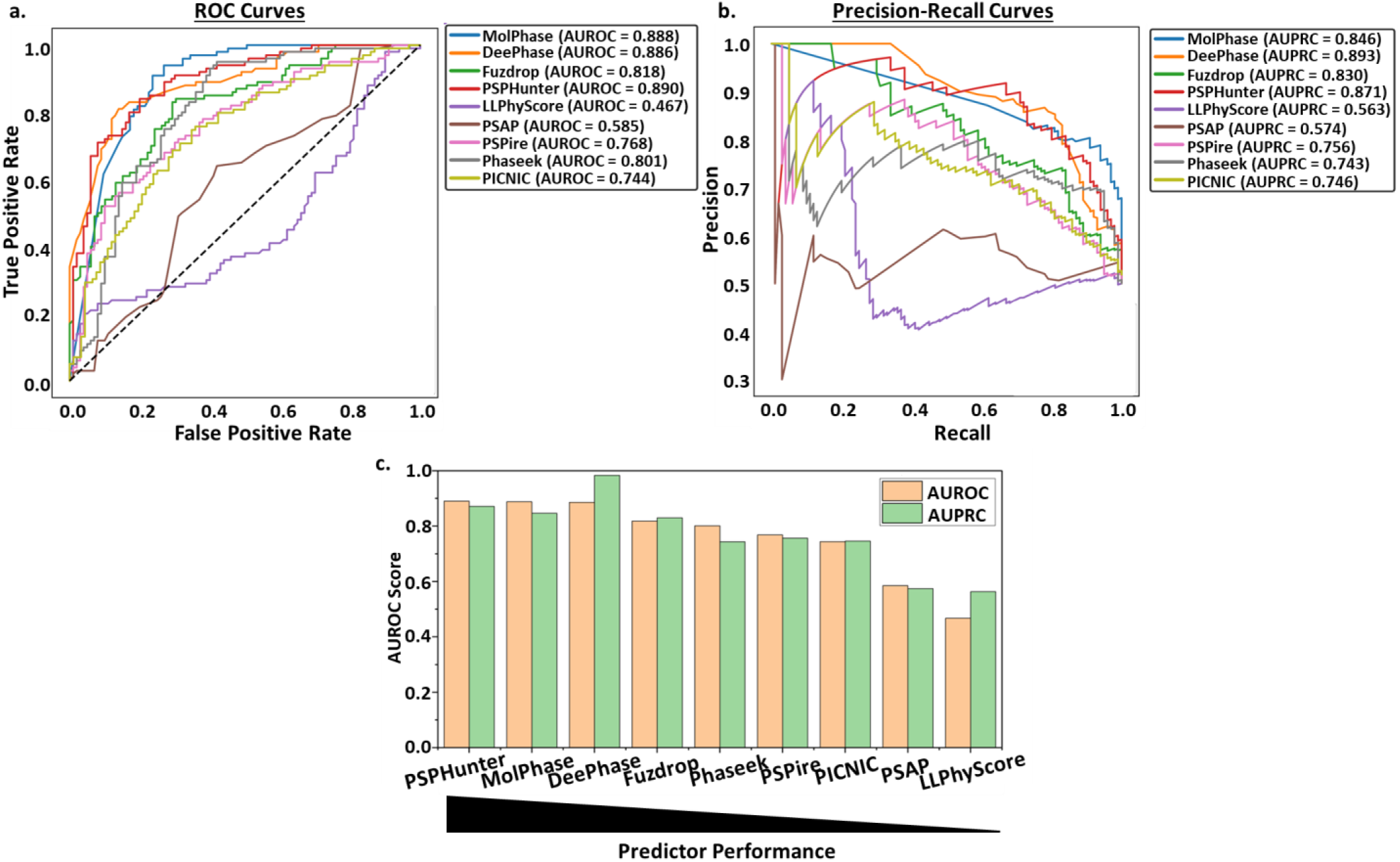
New ‘Benchmark’ datasets highlight the issues with current testing and training datasets. **a.** ROC and **b.** PRCs of all nine algorithms following prediction of the ‘Benchmark’ protein dataset. AUROC and AUPRC calculated for each. Black dashed line represents an AUROC of 0.5. **c.** Scores from a. and b. ranked from highest to lowest based on AUROC score.

LLPhyScore, PSAP, and PICNIC all showed the opposite trend, with increases in AUROC from the standard dataset. While suggesting that these predictors are being affected by incorrect labelling, their overall rank in comparison to others remain the lowest (Fig. 4c).

Overall, this exemplifies the importance of producing standardised benchmark datasets for both the training and testing of predictors. The currently used PDB and human proteome negative sets have inherent issues in their lack of protein diversity and likely inclusion of false negatives respectively. The inclusion of disordered negative proteins (e.g. DisProt) will aid in the removal of bias by increasing dataset diversity but will likely introduce false negatives similar to those seen with the human proteome. Therefore, the approach taken by Pintado-Grima et al. (2025) utilising a diverse negative dataset with stringent filtration provides the best balance of diversity with reducing type II errors.

### Mutation-level analysis exposes weaknesses in predictor robustness and adaptability

A growing area of interest in LLPS research is the contribution of single residues or small regions in the protein sequence in driving the phase separation process. Numerous PSPs have been identified as being driven by such regions and when these get mutated or removed, it severely inhibits or completely ablates the ability for those proteins to undergo LLPS (48–50). As aberrant phase separation has been linked to a variety of diseases (7–11), it is of interest to be able to predict the following: a) regions thought to drive phase separation, and b) the impact of mutating these regions on overall protein phase separation ability. While a growing number of algorithms are providing residue-level information involved in predictions (see Table 1), it is often stated that current predictors are unable to detect these changes reliably.

To investigate the ability of current second- and third-generation phase separation predictors to interpret the impact on LLPS caused by such changes in protein sequence, the *Mutations* dataset was produced. Alongside categorising LLPS+ proteins, LLPSDB v2.0 (51) also categorises experimentally established mutations in PSPs that can alter or completely disrupt its ability to undergo LLPS. The WT sequences for these mutations were collected and any with >40% similarity to those used in the predictor training datasets were removed. This returned a set of 12 proteins with 24 unique mutations sequences. For the purpose of this analysis, WT sequences represent the LLPS+ class, and the mutant sequences represent the LLPS- class.

As the class sizes for this dataset are significantly lower than those of the other testing datasets used, the AUROC and AUPRC must be interpreted carefully. With such a small dataset, these analysis parameters are particularly susceptible to even single incorrect predictions, causing large swings in the results. Additionally, the results will have limited resolution due to fewer datapoints causing coarse and less interpretable curves. Despite this, however, these metrics still provide good indicative insights into the predictors.

Across all predictors, the AUPRC scores (Fig. 5b) have reduced compared to the *Standard* dataset with three predictors having AUPRC below 0.5 (LLPhyScore, PSAP, and Phaseek) and only one predictor being above 0.7 (Deephase, AUPRC 0.893). In the context of this dataset, the LLPS+ class is 33% of the dataset meaning a random classifier would be expected to have an AUPRC of ∼0.33, a threshold of which all algorithms exceed except Phaseek which is just slightly under (0.321). This overall decrease and low levels of AUPRC show that the precision and recall in this situation is reduced, indicating all predictors (except DeePhase) struggle with correctly identifying the positives from the negatives.

**Figure 5.**
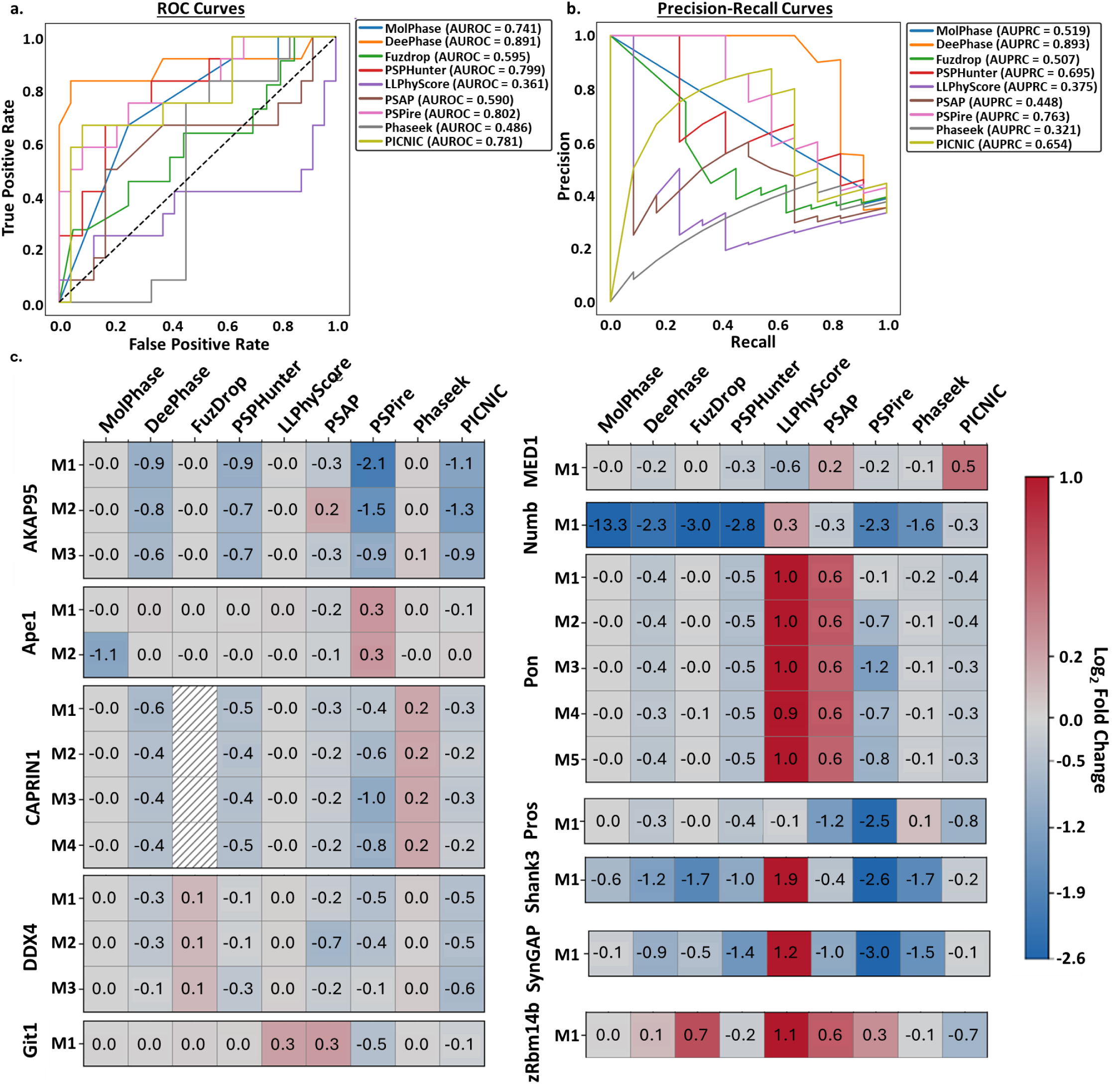
DeePhase, PSPHunter, PSPire, and PICNIC consistently detect LLPS propensity changes induced by small residue mutations. **a.** ROC and **b.** PRCs of all nine algorithms following prediction of the ‘Mutations’ protein dataset. AUROC and AUPRC calculated for each. Black dashed line represents an AUROC of 0.5. **c.** Log2 fold change calculated of prediction scores from the WT LLPS+ sequence to the mutant LLPS- variants. Red colours represent an increase in mutant LLPS propensity compared to WT. Blue colours represent a decrease in mutant LLPS propensity compared to WT.

Looking at the AUROC (Fig. 5a) shows us that top predictors have changed drastically. DeePhase remains the top performer in both metrics, showing minimal decreases in performance across all datasets assessed so far. However, the other two predictors making up the top three include PSPire and PSPHunter. As discussed previously, PSPire is a structure-informed predictor which has performed well in terms of AUROC but middling in comparative analysis for the other datasets. Alongside the other structure-informed predictor PICNIC, they have climbed the ranks when assessing the effects of mutations, suggesting including structural information in predictions may not have a large impact on overall prediction score but bolsters the ability to identify impacts on LLPS propensity caused by small mutations. The other top performer, PSPHunter, is an interesting algorithm as it has been designed with this specific purpose in mind – identifying phase-driving regions of proteins and the impacts of single mutations on propensity. For this study, the protein-level prediction module was used to enable calculation of analysis metrics and direct comparison to other algorithms, rather than the other purpose built-modules. Even in this instance, PSPHunter scored well, providing a high level of accurate ranking.

While it may be difficult for the predictors to directly give a binary ‘yes or no’ answer based on these small changes, they may be able to represent the change in LLPS ability as a decrease in propensity. For example, DeePhase identifies proteins with a score ≥0.5 as LLPS+ and those <0.5 as LLPS- and such, we would expect the WT sequence to be ≥0.5 and the mutant sequence to be <0.5. However, as detecting such small changes is a difficult classification task, the differences may be more in the lines of 0.85 for WT and 0.65 for the mutant. To assess this, the difference in LLPS prediction score between WT and mutant sequence was calculated as shown in Fig. 5c. Here, we can see three main outcomes for the predictors. The first is exemplified strongly by LLPhyScore and partially by PSAP in which an increase in LLPS propensity is often seen for the LLPS-mutants. This likely links to issues identified previously with these predictors in that they appear to be affected by bias and a lack of diversity in their training, and such have learned incorrect patterns in sequence features vs. LLPS propensity. The second type is shown well by MolPhase and Phaseek which are unable to detect changes caused by the LLPS- mutations compared to the WT sequence. For these predictors, residue-level changes have limited impact on their overall score and as such they are better suited for protein-level investigations rather than residue-level. The final type is represented by PSPire and PSPHunter in which quite large decreases in score are often reported. PSPHunter is the only algorithm that detected a decrease in almost all mutations and never reported an increase, showing its built-for-purpose algorithm is working optimally.

Overall, only a small subset of predictors were able to accurately detect decreases in LLPS propensity caused by small mutations and even then, are better represented as a fold change from WT rather than a binary result. PSPHunter is a built-for-purpose algorithm providing highly accurate predictions for small mutations in its protein-level predictions with additional modules allowing for further investigation at the region- and residue-level. Interestingly, structure-informed predictors also excel in this area compared to their performance with other datasets, suggesting structural information may aid in the identification of small regions and residues rather than a protein-level overall prediction. Interestingly, there appears to be little crossover in key features of the top performing predictors that are unique from those with weaker performance. Both sequence-based predictors, DeePhase and PSPHunter, do employ word2vec embeddings, treating residues and domains as ‘words’ in vectors which has been used for many types of protein prediction (63–66), potentially explaining their success. A more in-depth analysis is beyond the scope of this study but could provide key insights into which machine learning or protein-level features are unique to accurate mutation effect prediction and if structure could play a part in improving it further.

### Reduced Accuracy on Viral Proteins Suggests a Distinct Molecular Signature Not Captured by Current Models

An area of LLPS research that has been recently increasing in popularity is that of LLPS in viral replication cycles. Many viruses form replication centres in the cell by large-scale remodelling of cellular membranous organelles such as the Golgi apparatus (67), endoplasmic reticulum (68–71), and lysosomes (72,73). This allows for the formation of a ‘viral factory’ in which key cellular and virus components are concentrated to allow efficient replication and virion production (74). However, recently, an increasing number of viruses have been shown to induce the formation of liquid-organelles in order to build their viral factories, driven by viral proteins via the process of LLPS (75–80). This has had a large impact on the understanding of virus replication cycles and how viruses are able to hijack cells successfully. It is therefore important to be able to accurately and rapidly screen viral proteins predicted to phase separate to accelerate research into virus-based LLPS. However, viral proteins are underrepresented in many databases used to collate training datasets (28) and often do not have homologues in other proteomes (29) thus predictors are unlikely to have learned any virus-specific patterns.

To investigate how well predictors can handle viral proteins, a *Virus* dataset was produced. The LLPS+ class was collected from BAV-LLPS-DB, a newly produced LLPS database dedicated to PSPs from bacteria, archaea, and viruses (52). For curation of the LLPS- dataset, the *Standard* approach was taken in which virus-specific proteins with high-resolution X-ray structures were chosen. While the issues with using PDB-only LLPS-sequences have been discussed thoroughly here, it was chosen for the *Virus* dataset for two main reasons. First, it ensures the dataset is as similar to the *Standard* as possible, providing the clearest comparison to determine if the predictors face issues with virus-specific sequences rather than a mixed set. Second, due to the limited number of entries in the LLPS+ class and the need to condense the LLPS- dataset to match, the PDB dataset currently provides the highest likelihood of containing fewer false negative proteins.

Across all predictors tested, the AUROC and AUPRC of each (Fig. 6a, b) decreased compared to that of the *Standard* dataset (Fig. 2a, b). The largest reduction was with Phaseek with the AUROC decreasing by 0.329 from 0.911 to 0.582. DeePhase, while still performing well (0.866) dropped in the rankings (Fig. 6c) with a decrease in AUROC score of 0.115. Despite this, however the top three algorithms still remained as MolPhase, Deephase, and Fuzdrop, showing their high accuracy across all datasets, except for the *Mutation* set.

**Figure 6.**
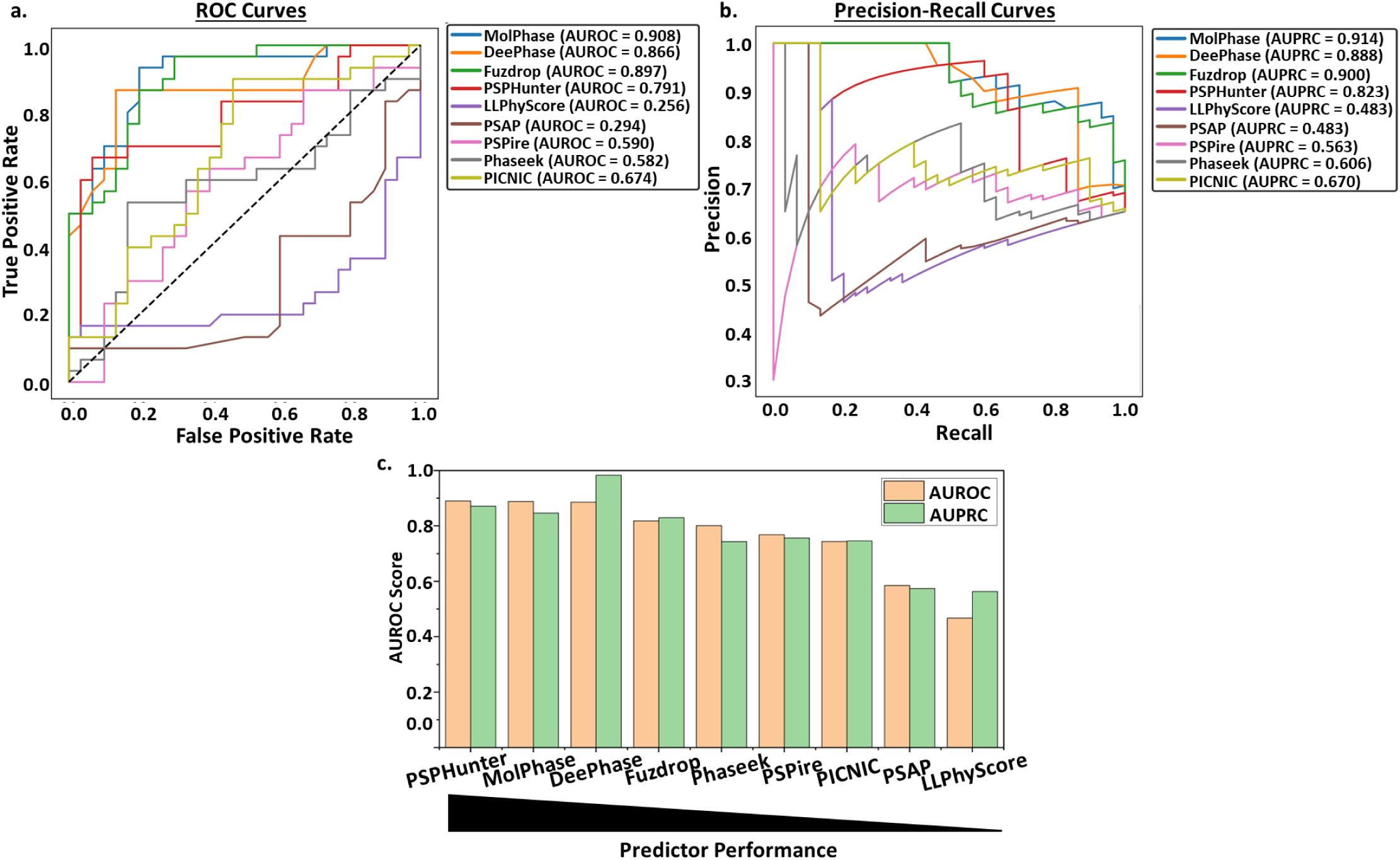
LLPS predictors are less confident assessing virus-specific proteins. **a.** ROC and **b.** PRCs of all nine algorithms following prediction of the ‘Virus’ protein dataset. AUROC and AUPRC calculated for each. Black dashed line represents an AUROC of 0.5. **c.** Algorithms ranked from best to worst performance based on AUROC score.

Overall, this highlights that viral proteins likely have a somewhat different ‘molecular language’ to that of proteins from different organisms. Thus, the training of these predictors, often either on a virus-underrepresenting database or on the human proteome alone, does not capture virus LLPS as accurately. To further investigate this, a dataset should be created of virus-only proteins and utilised to train high-performing prediction algorithms either alone or alongside standard datasets to understand if these patterns can be used to ascertain more accurate prediction scores.

## Conclusion

As the research area of LLPS in biological systems continues to grow and merge with other disciplines, the need for accurate and user-friendly LLPS predictors gets larger. This has led to the development of a wide array of second-generation machine learning algorithms designed to learn the ‘molecular language’ of proteins and identify features characteristic of PSPs. However, with the variety of predictors available, each utilising different protein characteristics from sequences or structures, and employing various machine or deep learning approaches, it is difficult to decide on which to use. This study provides an unbiased comparative analysis of nine of the currently available second- and third-generation predictors across a comprehensive collection of datasets. An overview of the performance of each predictor can be seen in Fig. 7.

**Figure 7.**
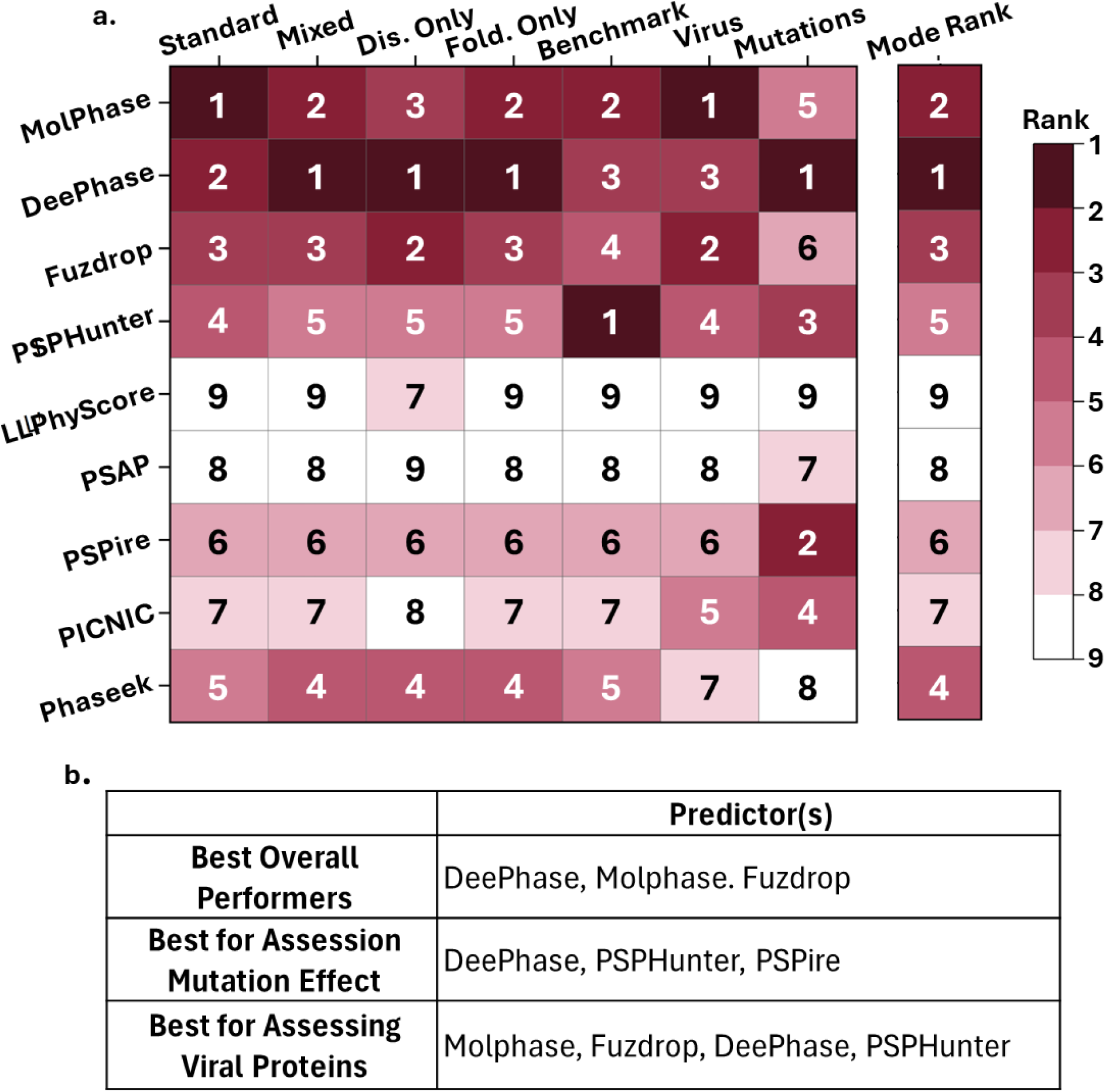
Overall performance of nine LLPS predictors. **a.** Heatmap showing the “rank” of all nine algorithm across each dataset based on highest (1st) to lowest (9th) AUROC score. Overall rank was determined by collecting the modal value for each predictor. Dark red colours represent higher scoring predictors and light red to white represent the lower scoring predictors. **b.** Table showing the best performing algorithms for specific usage situations.

DeePhase provided the best overall performance in all datasets, never falling below the top three performers (Fig. 7a), indicating it provides accurate predictions across a diverse range of proteins. MolPhase also performs similarly to DeePhase across the board, only faltering when assessing the impact of small mutations on protein-level LLPS propensity. MolPhase does, however, provide a larger amount of insight into residue-level details for each metric considered, as outlined in Table 1. Fuzdrop is the third best ranked predictor, however it is outperformed on every dataset by another, though it does beneficially provide information on specific regions in the protein that are predicted to promote LLPS or aggregation. Therefore, for the purposes of screening proteins for LLPS propensity, DeePhase or MolPhase provide the most accurate results.

As many researchers are interested in the investigation of residues or regions that are key for driving LLPS and such can be mutated to abolish or disrupt it, it is important to identify predictors useful in this specific situation. Somewhat surprisingly, DeePhase was the best performing in this dataset in terms of AUROC despite not being specifically designed for this purpose. Even when considering the evaluation metric of fold change rather than a binary ‘yes or no’ DeePhase was able to report a decrease in most of the mutant sequences compared to WT. However, here it was outperformed by PSPHunter, the built-for-purpose predictor for identifying deleterious mutations and phase-driving regions. Interestingly, two other high performers here were the two third-generation structure-informed predictors PSPire and PICNIC which had otherwise performed middling in the other datasets. This suggests that while structural data may not be as important as sequence data for prediction of an overall protein-score, it can provide good insight into the effect of small mutations on overall propensity. Therefore, to investigate the impact of small mutations on a protein and their phase-driving regions, a mixed-approach utilising multiple predictors will likely give the most comprehensive insight, specifically DeePhase, PSPHunter, and PSPire.

Another area of growing research regarding LLPS is the involvement of it in viral replication processes. LLPS is being implicated as a key process in numerous viruses and as such, the need to be able to predict which virus proteins are likely to undergo this is increasing. However, many of these algorithms are trained on datasets in which viral proteins are underrepresented and so any LLPS-inducing patterns unique to viral proteins is likely being missed. This study has identified that this is a likely bias in many of the predictors currently existing but with the best performer for these sequences being MolPhase which showed limited AUROC differences in comparison.

Throughout this study, it has been well-documented the issues that currently persist in the training and testing datasets used for many predictors. They are currently showing bias towards the outdated view that disorder guarantees LLPS and majority-folded proteins cannot undergo LLPS. It is imperative to create both LLPS+ and LLPS- datasets that represent the wide variety of proteins in each class. One method of doing this utilised in this study is shown by Pintado-Grima et al. (2025) in which proteins from well-structured and disordered protein databases were used in the LLPS- dataset and were strictly filtered to remove as many false negatives as possible. The use of such standardised benchmark datasets could then lead to a new generation of LLPS predictors with the best prediction accuracy yet.

## Additional Files

Additional_File_1.xlsx – **Individual protein sequences and corresponding LLPS predictor scores across datasets.** Spreadsheet containing the individual FASTA protein sequences utilised in each dataset and the individual scores calculated by each algorithm for each dataset used for data analysis.

## Declarations

### Ethics approval and consent to participate

Not applicable

### Consent for publication

Not applicable

### Availability of data and materials

All data generated and analysed during this study are included in this published article and its supplementary information files.

### Authors’ contributions

K.E.G performed data curation, analysis, methodology, data visualisation, and original draft writing. J.N.B performed funding acquisition, and manuscript review and editing. All authors have read and agreed to the published version of the manuscript.

## Acknowledgements

Not applicable

## Authors’ information (optional)

Not applicable

